# Street RABV induces the cholinergic anti-inflammatory pathway in human monocyte-derived macrophages by binding to nAChr ɑ7, and upregulates the M2-c marker CD163

**DOI:** 10.1101/2021.01.05.425407

**Authors:** C.W.E. Embregts, L. Begeman, C.J. Voesenek, B.E.E. Martina, M.P.G. Koopmans, T. Kuiken, C.H. GeurtsvanKessel

## Abstract

Rabies virus (RABV) is able to reach the central nervous system (CNS) without triggering a strong immune response, using multiple mechanisms to evade and suppress the host immune system. After infection via a bite or scratch from a rabid animal, RABV comes into contact with macrophages, which are the first antigen-presenting cells (APCs) that are recruited to the area and play an essential role in the onset of a specific immune response. It is poorly understood how RABV affects macrophages, and if the interaction contributes to the observed immune suppression. This study was undertaken to characterize the interactions between RABV and human monocyte-derived macrophages (MDMs). We showed that street RABV does not replicate in human MDMs. Using a recombinant trimeric RABV glycoprotein (RABV-tG) we showed binding to the nicotinic acetylcholine receptor alpha 7 (nAChr ɑ7) on MDMs, and confirmed the specificity using the nAChr ɑ7 antagonist alpha-bungarotoxin (ɑ-BTX). We found that this binding induced the cholinergic anti-inflammatory pathway (CAP), characterized by a significant decrease in tumor necrosis factor ɑ (TNF-ɑ) upon LPS challenge. Using confocal microscopy we found that induction of the CAP is associated with significant cytoplasmic retention of nuclear factor κB (NF-κB). Co-cultures of human MDMs exposed to street RABV and autologous T cells further revealed that the observed suppression of MDMs affects their function as T cell activators as well, as we found a significant decrease in proliferation of CD8^+^ T cells. Lastly, using flow cytometric analysis we observed a significant increase in expression of CD163, hinting that street RABV is able to polarize macrophages towards a M2-c anti-inflammatory phenotype. Taken together, these results show that street RABV is capable of inducing an anti-inflammatory state in human macrophages, which affects T cell proliferation.

**Author summary:** Rabies virus (RABV) is transmitted by a bite or a scratch from an infected animal. Infection leads to a lethal encephalitis and once clinical symptoms occur, there is no effective treatment available. The virus is able to travel from the initial site of infection to the central nervous system without triggering a strong immune response, using multiple mechanisms to evade and suppress the immune system. Up to present it is unclear when and where this immunosuppression is initiated, and if local immune cells are involved as well. Understanding the complete mechanisms of immunosuppression by RABV is essential for the development and improvement of effective post-exposure treatments. In this paper we studied if RABV is able to suppress human primary macrophages as these will be the first antigen-presenting cells that are recruited to the site of infection, and are known to be important in initiating an efficient immune response. We show that RABV is able to bind, but not infect, human macrophages. Binding induces an anti-inflammatory pathway, which leads to limited T cell proliferation and directs macrophages towards and anti-inflammatory state. These results show that RABV-macrophage interactions might indeed be one of the early steps in the onset of RABV-induced immunosuppression.

## Introduction

Rabies is a zoonotic viral encephalitis responsible for 60 000 reported human deaths annually, although the true burden is suspected to be much higher (Hampson et al., 2015). The disease is caused by members of the genus *Lyssavirus*, -ssRNA viruses of the *Rhabdoviridae* family. While multiple lyssaviruses have been reported to cause rabies in humans, over 99 percent of all reported human cases are caused by dogs infected with rabies virus (RABV) (WHO, 2018). Nevertheless, in North and South America bats are the main source of human infections, mostly caused by silver-haired bat rabies virus (SHBRV) (Dietzschold et al., 2000).

While effective pre- and post-exposure prophylaxis is available, no treatment options are currently available once clinical symptoms occur. This makes rabies the deadliest zoonosis with a case-fatality rate approaching 100%. The virus is able to reach the CNS without triggering a strong local immune response (Schnell et al., 2010; Yamaoka et al., 2013); neither is a systemic immune response induced, as indicated by the low or absent neutralizing antibodies at the end-stage of disease of most patients (Kasempimolporn et al., 1991; Noah et al., 1998). This lack of immune activation is caused by limited viral replication at the inoculation site (Charlton et al., 1997; Murphy and Bauer, 1974), as well as by active immune evasion and suppression, as extensively reviewed (Katz et al., 2017; Scott and Nel, 2016). Previous research identified major immune evasive mechanisms, including the blocking of RIG-I activation by the nucleoprotein (N) (Masatani et al., 2013, 2011, 2010) and the IFN signal transduction pathway by the phosphoprotein (P) (Brzózka et al., 2006, 2005; Vidy et al., 2005).

To date, the majority of research on immunosuppressive mechanisms focused on the response in neurons and the CNS. While the absence of an effective local immune response after infection with RABV has been described (Yamaoka et al., 2013), it is not clear how exactly RABV affects local immune cells at the site of infection. Suppression of local immune cells may be responsible for the lack of immune response in later stages of diseases, hence increasing the knowledge on interactions between RABV and local immune cells is indispensable in the development of effective and targeted treatment strategies. Given this importance, we aimed to investigate the functional effects of street RABV on macrophages. Macrophages are versatile innate immune cells and play an essential role in the uptake and clearance of pathogens, and initiation of a strong local response by producing pro-inflammatory cytokines, chemokines and nitric oxide (NO) (Koh and Dipietro, 2019; Mosser and Edwards, 2008). After being bitten or scratched by a rabid animal, RABV first encounters resident macrophages in skin and muscle tissue, after which the inflicted physical damage and the presence of bacteria and other microbes in the wound area induce a strong influx of immune cells (Brancato and Albina, 2011; Daley et al., 2010). Large numbers of infiltrated macrophages have been found in peripheral sites after inoculation with RABV as well (Charlton and Casey, 1981); however, the interactions between RABV and macrophages have been studied incompletely. The few studies focusing on the direct effects of RABV on macrophages showed that various RABV strains were able to induce NO production and CXCL10 expression in RAW264 murine macrophages (Nakamichi et al., 2004), and induced macrophage apoptosis through involvement of various caspases (Kip et al., 2017). In the aforementioned papers, lab-adapted or attenuated RABV strains and murine macrophage-like cell lines were used, therefore the effects of street RABV strains on primary macrophages remain unstudied.

Macrophages have a high functional plasticity. Upon encountering pathogens, and in combination with cytokines produced by resident cells, macrophages polarize towards diverse and distinct functional phenotypes, as are normally grouped into classically activated or inflammatory M1 phenotypes, and alternatively activated or anti-inflammatory M2 phenotypes (Stout et al., 2005). A typical type 1/Th1 immune response, associated with M1 macrophages, aids the elimination of intracellular pathogens while a typical type 2/Th2 response, associated with M2 macrophages, will reduce viral clearance. Shifting macrophage polarization towards a phenotype beneficial for the virus is a strategy known for many viruses, as extensively reviewed (Sang et al., 2015). Studies on the effects of street RABV strains on macrophage polarization are lacking.

Nicotinic acetylcholine receptors (nAChr) are a family of ligand-gated pentameric ion channels with a variety of pharmacological functions. The nAChr ɑ7 is a homopentameric nAChr that consists of five ɑ7 subunits and is involved in the cholinergic anti-inflammatory pathway (CAP), characterized by a decreased inflammatory response of macrophages upon binding of a typical ligand (nicotine, acetylcholine) to nAChr ɑ7 (Wang et al., 2003). RABV glycoprotein (RABV-G) can bind to nAChr ɑ7, but functional effects of this binding have not yet been studied (Kim et al., 2010). We hypothesize that binding to the nAChr ɑ7 leads to suppression of the inflammatory response of macrophages.

In the present study we showed that a street RABV strain does not replicate in human monocyte-derived macrophages (MDMs). We confirmed specific binding of RABV-G to nAChr ɑ7 on human MDMs and, for the first time, showed that a viral protein (RABV-G) was able to induce the cholinergic anti-inflammatory pathway (CAP) in MDMs. We found that the induction of the CAP was related to cytoplasmic retention of NF-κB in RABV-exposed MDMs, and that exposure of MDMs with RABV decreased proliferation of autologous T cells *in vitro*. In parallel, exposure of MDMs to RABV did not induce typical M1 phenotypic markers, but instead upregulated the M2-c phenotypic marker CD163, hinting at the possibility that RABV is able to shift macrophage polarization towards an anti-inflammatory M2-like phenotype.

## Methods

### Virus

A street rabies virus (RABV) strain known for its capability to cause human infections in North America, the silver-haired bat rabies virus (SHBRV), was used. The virus was propagated in the human neuroblastoma cell line SK-N-SH in Eagle’s Minimum Essential Medium (EMEM) with Earle’s Balanced Salt Solution (EBSS) (Lonza), supplemented with 10% (v/v) Fetal Calf Serum (FCS), 100 U penicillin (Gibco), 100 mg/mL streptomycin (Gibco), 2 mM L-glutamine (Gibco), 1% nonessential amino acids (Lonza), 1 mM sodium pyruvate (Gibco) and 1.5 mg/mL sodium bicarbonate (Gibco). Virus titrations were performed by the median tissue culture infective dose (TCID_50_) endpoint dilution method of Reed and Muench (Reed and Muench, 1938) using the mouse neuroblastoma cell line MNA. MNA cells were cultured in Dulbecco’s Modified Eagle Medium (DMEM, Gibco) supplemented with 10% FCS, 100U penicillin, 100 mg/mL streptomycin, 2 mM L-glutamine and 1 mM sodium pyruvate (Gibco). SK-N-SH and MNA Cells were maintained at 37 °C with 5% CO_2_.

### Primary cell isolation and macrophage maturation

Peripheral blood mononuclear cells (PBMCs) were isolated from buffy coats obtained from healthy, non-smoking and non-rabies vaccinated individuals (Sanquin). PBMCs were obtained by density centrifugation using Ficoll Paque PLUS (GE Healthcare). Monocytes and T cells were obtained from PBMC fractions by magnetic associated cell sorting using CD14^+^ and CD3^+^ beads, respectively, following manufacturers guidelines (Miltenyi Biotec). Purity of the sortings was confirmed by flow cytometry using a BD Lyric flow cytometer (BD Biosciences).

Monocytes were seeded at a density of 100 000 cells per well in 96-well plates and were maturated for six days in Roswel Park Memorial Institute-1640 (RPMI-1640) medium containing 10% pooled human serum (Sanquin), 1% (v/v) GlutaMAX (Gibco) and 20 ng/mL monocyte colony-stimulating factor (M-CSF, R&D Systems). Cells were maintained at 37 °C with 5% CO_2_ and the medium was replaced on day 2 and day 4.

### Infection of human monocyte-derived macrophages

Mature human monocyte-derived macrophages, obtained on day six of culture, were exposed to RABV for one hour in serum-free medium using a MOI of 0.1, 1, 10 or 50, after which the cells were washed once and were incubated in complete medium. At 8, 24, 48, and 72 hours post-infection, cell culture supernatants were harvested for virus titration and cells were fixed in 80 % acetone for immunofluorescent detection of the nucleoprotein using a mouse FITC-labeled RABV-N antibody (Fujirebio). Nuclear counterstaining with Hoechst 33342 (Sigma Aldrich) was included before image acquisition using a Zeiss LSM700 confocal laser scanning microscope. Virus titrations were performed by the TCID_50_ endpoint dilutions method of Reed and Muench using the mouse neuroblastoma cell line MNA, as described before. A total of six donors were included for the MOIs 0.1 and 1, and two donors for the MOIs of 10 and 50. Two wells were infected for every condition and all supernatants were titrated in triplicate. Infection of the highly susceptible neuroblastoma cell lines SK-N-SH and MNA cells were included as positive controls for virus infectiousness.

### rRABV-tG binding assay

The ability of the RABV glycoprotein to bind to human monocyte-derived macrophages was investigated using a recombinantly expressed trimeric form of RABV-G (rRABV-tG, described in Koraka et al., 2014). rRABV-tG was labeled with a 0.1 mg/mL solution of FITC (Sigma Aldrich) in 0.5M bicarbonate buffer (pH 9.5) for one hour under constant stirring, after which unbound FITC was removed by overnight dialysis in PBS. Mature macrophages were incubated with 50 μg/mL FITC-labeled rRABV-tG for 30 minutes at 4 °C. In parallel, macrophages were incubated with ATTO-633 conjugated alpha-bungarotoxin (ɑ-BTX; Alomone labs) or with an antibody against the nicotinic acetylcholine receptor ɑ7 (nAChr ɑ7; Alomone Labs) followed by an Alexa594 goat-anti-rabbit conjugate (Invitrogen). Nuclear counterstain with Hoechst 33342 (Sigma) was included before cells were imaged using a Zeiss LSM700 confocal laser scanning microscope.

For quantification of rRABV-tG binding, mature macrophages (n=6 individual donors) were detached using Accutase (Merck Millipore), blocked with 10% pooled human serum in PBS for 30 minutes and incubated with various concentrations of rRABV-tG (0-100 μg/mL) for 30 minutes at 4 °C. To verify the binding of rRABV-tG to the nAChr-ɑ7, macrophages were incubated with various concentrations of the nAChr ɑ7 antagonist alpha-bungarotoxin (ɑ-BTX, 0-50 ug/mL) for 15 minutes before addition of rRABV-tG. After two washes in FACS buffer, binding of rRABV-tG, as well as the expression of nAChr ɑ7 was quantified using a BD Lyric flow cytometer (BD Biosciences). Data were analyzed using FlowJo V10.6.2.

### Macrophage stimulation and TNF-ɑ cytokine assay

The anti-inflammatory effect of RABV on macrophages was studied using a lipopolysaccharide (LPS) stimulation. LPS is an outer membrane component of Gram-negative bacteria and is known to induce a strong inflammatory response. Briefly, mature macrophages (n=6–9 donors) were exposed to RABV (MOI of 10, 25 or 50) for one hour and subsequently with LPS (100 ng/mL) for six hours. Treatment with acetylcholine (1 μg/mL) was used as a positive control for induction of the cholinergic anti-inflammatory pathway. Wells without LPS challenge, or LPS challenge without pre-exposure to infectious RABV or acetylcholine, were included as negative and positive control, respectively. To confirm the binding of RABV to nAChr ɑ7, the specific antagonist alpha-bungarotoxin (ɑ-BTX, 2 µg/mL) was used as a pre-treatment before cells were exposed to RABV or acetylcholine, and subsequently to LPS (Fig 3A). All conditions were tested in triplicate wells. Supernatants were collected after six hours of stimulation with LPS and TNF-ɑ concentrations were determined using the Legendplex TNF-ɑ cytometric bead assay (BioLegend). Bead analysis was performed by a BD FACS Lyric flow cytometer and the data was analyzed using FlowJo V10.6.2. Relative TNF-ɑ reduction per individual donor was obtained by normalizing MFI’s against the TNF-ɑ production of the corresponding wells stimulated with LPS only.

### Quantification of NF-κB nuclear translocation and retention

Macrophages (n=6 donors) were exposed to RABV (MOI of 50) or acetylcholine (1 μg/mL) and afterwards challenged for six hours with LPS (100 ng/mL) as described above. Wells were washed twice with PBS, fixed with 4% paraformaldehyde (PFA) in PBS for 15 minutes and permeabilized using PBS 0.1% Triton X-100 for 15 minutes. After blocking for 30 minutes with 2% FCS in PBS, wells were stained with a monoclonal antibody against the NF-κB the p65 subunit (Santa Cruz) and goat-anti-mouse Alexa488 (Invitrogen). Nuclei were counterstained with Hoechst 33342 (Sigma Aldrich) and wells were imaged using a Zeiss LSM700 confocal laser scanning microscope.

An ImageJ macro was designed to perform automatic and user-independent quantification of nuclear and cytoplasmic NF-κB. Briefly, z-stacks were projected into single images by summing the intensity of all slices. Nuclear regions were defined by the Hoechst 33342 staining and a binary mask was created using an intensity threshold. Total NF-κB was detected by applying the Li intensity threshold on the Alexa488 channel, after which the signal was converted into a binary mask. In order to obtain the cytoplasmic NF-κB, the binary mask containing the nuclei was subtracted from the total NF-κB channel. The area and mean NF-κB intensity were measured in the original z-projection. Lastly, total NF-κB expression was determined by area * intensity of both the nuclear and cytoplasmic region and the average nuclear/cytoplasmic ratios were determined for each image. At least three high-magnification fields were imaged per well, and nuclear/cytoplasmic ratios were determined for six donors in total.

### T cell co-culture

Macrophages (n=6 donors) treated with RABV (MOI of 50) for one hour were washed twice with medium (RPMI supplemented with10% FCS, 100U penicillin, 100 mg/mL streptomycin and 2 mM L-glutamine) before autologous T cells were added. CD3^+^ T cells were stained with 2 μM carboxyfluorescein succinimidyl ester (CFSE) as described before (Piazzon et al., 2015), were activated with 2.5 μg/mL ɑ-CD3 (Invitrogen) and ɑ-CD28 (Invitrogen) and were co-cultured with macrophages in a 1:1 ratio. Non-stimulated T cells, as well as stimulated T cells cultured without macrophages, were taken along as controls for each donor. Cells were cultured in 3.5 days after which proliferation was determined by flow cytometry. To this end, T cells were washed twice with PBS, stained with the fixable viability dye ZombieViolet (Biolegend), fixed with 4% PFA for 15 minutes and subsequently stained for 30 minutes with anti-CD4-APC and anti-CD8-BV605 (Biolegend) in FACS buffer. Flow cytometry analysis was performed using a BD FACS Lyric flow cytometer and the data were analyzed using FlowJo V10.6.2.

### Macrophage maturation and polarization

Mature macrophages (n = 6 donors) were stimulated for 48 hours with complete medium containing IFN-γ (20 ng/mL, R&D Systems) and LPS (100 ng/mL, Sigma Aldrich), with IL-4 (20 ng/mL, R&D Systems), or with IL-1 (20 ng/mL R&D Systems), to induce the M1, M2a or M2c phenotype, respectively. To investigate the effect of rabies virus on macrophage polarization, macrophages were stimulated with complete medium containing RABV (MOI of 10). Expression of macrophage phenotypical markers was investigated by flow cytometry. Non-polarized macrophages, cultured for 48 hours with complete medium without additional cytokines, were taken along as controls for each donor.

Macrophages were dissociated from the wells using Accutase (Merck Millipore) and were washed twice with PBS before staining for 30 minutes with the fixable viability dye ZombieViolet (Biolegend). Cells were fixed with 4 % PFA for 15 minutes, and after Fc receptor blocking with Human TruStain FcX (Biolegend), cells were stained with the following antibodies in FACS buffer (PBS with 2 % fetal calf serum, 0.2 mM EDTA, 0.01% sodium azide): anti-CD80-FITC, anti-HLA-DR-APC-Cy7, anti-PD-L1-APC, anti-CD163-PE, anti-CD200R-PE-Cy7 and anti-CD206 (BV786) (All Biolegend). Mean fluorescent intensities were quantified by flow cytometry using a BD Lyric flow cytometer (BD Biosciences) and the data were analyzed using FlowJo V10.6.2.

### Statistical analysis

All statistical analyses were performed using SPSS Statistics 25 (IBM).

For the rRABV-tG binding assay, the TNF-ɑ cytokine assay and the polarization study, One-Way ANOVAs were performed followed by Tukey’s post-hoc tests to test significant differences between treatments. For the NF-κB assay and the proliferation assay, paired sample T-tests were performed to determine significant differences. *P* values < 0.05 were considered significant and all results were expressed as mean ± SEM.

## Results

### Street RABV does not replicate in human monocyte-derived macrophages

To determine the immunomodulatory effects of RABV-macrophage interactions, we first investigated if exposure to street RABV leads to a productive infection in human MDMs, measured by an increase in virus over time. To this end, mature MDMs were exposed to street RABV in a MOI of 0.1, 1, 10 or 50, and at 8, 24, 48, and 72 hours post-infection (hpi) the presence of virus was examined by fluorescent microscopy and virus titration. The highly susceptible human neuroblastoma cell lines MNA and SK-N-SH were taken along as positive controls. RABV-N staining was detected from 24 hpi onwards in the MNA and SK-N-SH cells, but remained absent throughout all time-points in the human MDMs (Fig 1A). MNA and SK-N-SH cells exposed to a MOI of 1 reached a plateau phase around 48 hpi, with titers between 10^6.17^ and 10^6.50^ TCID_50_/mL, and cells inoculated with a MOI of 0.1 reached this phase around 72 hpi (Fig 1B). While a low positive signal (10^1^ TCID_50_/mL) was detected at 24 hpi in human MDMs that were exposed to a MOI of 1, no increase was observed at 48 and 72 hpi, or in MDMs infected with a higher MOI This lack of increase of present virus, as well as the absence of RABV-N protein at all tested time points, shows that inoculation with a street strain of RABV does not lead to a productive infection of mature human MDMs.

**Fig 1.**
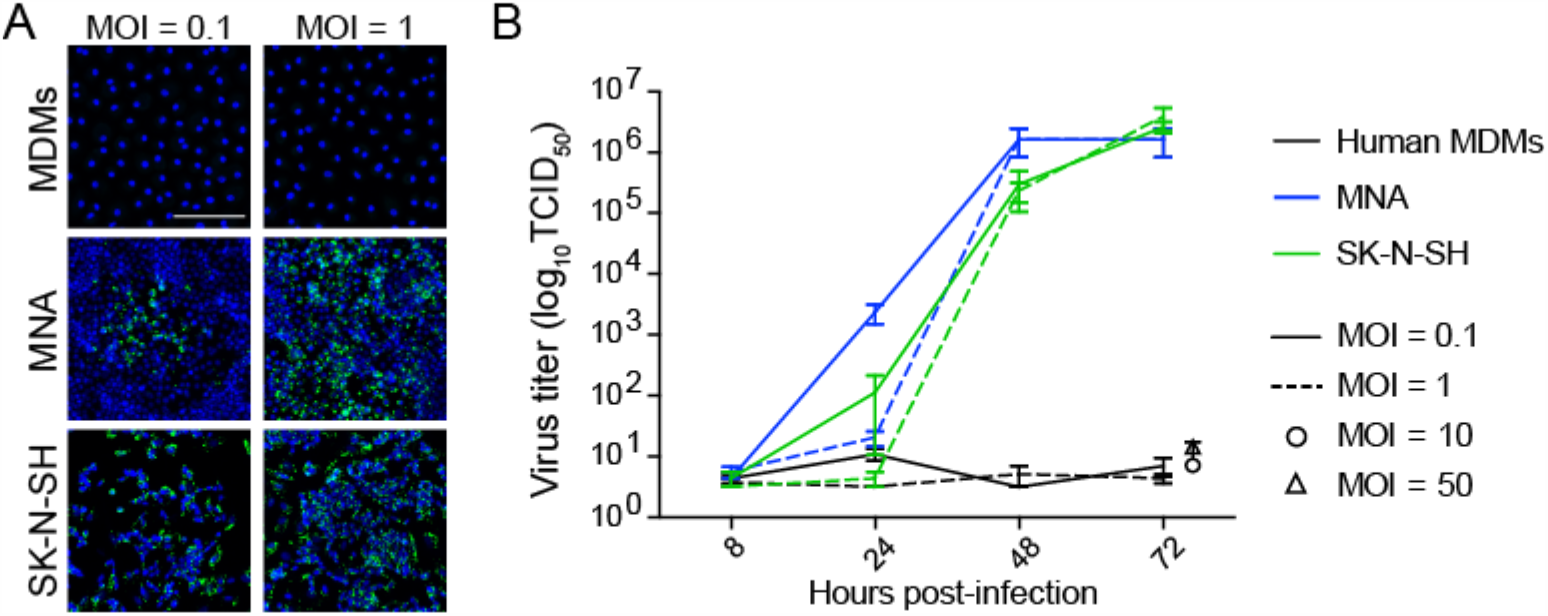
Qualitative and quantitative assessment of the ability of RABV to infect mature human monocyte-derived macrophages. Human monocyte-derived macrophages (MDMs) were exposed to street RABV at a MOI of 0.1, 1, 10 or 50, and at 72 hours post-infection (hpi) cells were fixed and stained with an antibody against the RABV-N protein (A). The white scalebar represents 500 μM and all images were acquired at the same magnification. In parallel, supernatants of 8, 24, 48 and 72 hpi were titrated on MNA cells (B). Bars represent the mean ± SEM of two-six donors; for each condition, two wells were infected per condition and all supernatants were titrated in triplicate. Highly susceptible MNA and SK-N-SH cells were taken along as positive controls.

### RABV glycoprotein binds to nAChr ɑ7 on human monocyte-derived macrophages

As a first step in testing if RABV is able to induce the cholinergic anti-inflammatory pathway (CAP) in human macrophages, we investigated the ability of RABV-G to bind to nicotine acetylcholine receptor alpha-7 subunit (nAChr ɑ7) on human MDMs. Binding was examined by both flow cytometry and confocal microscopy, using FITC-labelled recombinant trimeric RABV-G (rRABV-G). First, expression of nAChr ɑ7 was examined on mature MDMs by confocal microscopy (Fig 2A) and flow cytometry (Fig 2B), confirming a strong expression of this receptor. In parallel, investigation on the binding of nAChr ɑ7 antagonist alpha-bungarotoxin (ɑ-BTX) and the FITC-labeled rRABV-tG by confocal microscopy, showed that both ɑ-BTX and rRABV-tG could efficiently bind to human MDMs (Fig 2A). Next, macrophages were incubated with various concentrations of rRABV-tG, and flow cytometric analysis revealed a concentration-dependent binding to the MDM surface (Fig 2C). In order to confirm that rRABV-tG binds to nAChr ɑ7, a pre-treatment with ɑ-BTX was used. Although no significant inhibition of rRABV-tG binding was observed at any of the tested ɑ-BTX concentrations (0-50 µg/mL), binding showed a decreasing trend with increasing concentrations of ɑ-BTX (Fig 2C). This confirms that ɑ-BTX and rRABV-tG compete for the same receptor and that indeed rRABV-tG binds to nAChr ɑ7 on human MDMs.

**Fig 2.**
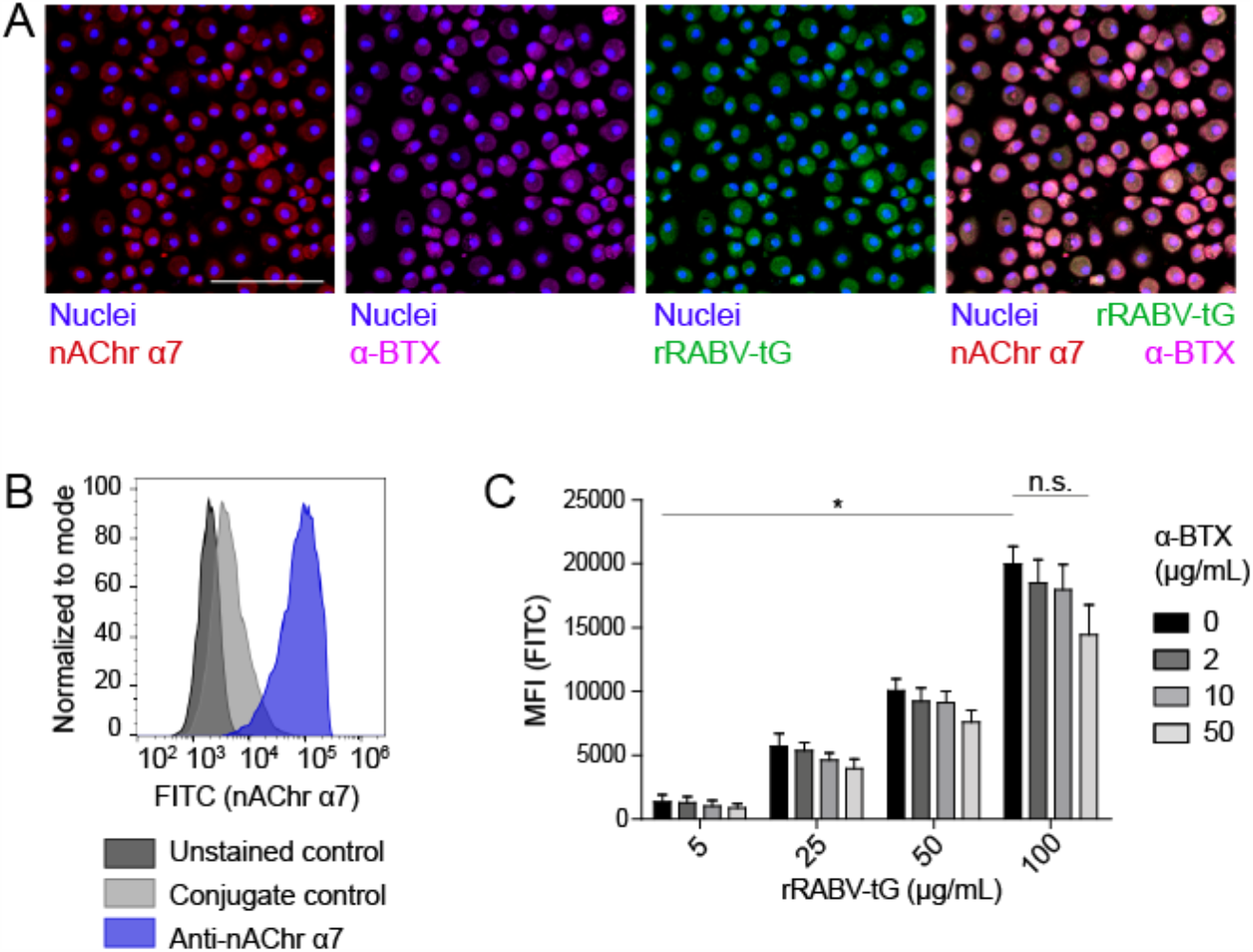
Presence of nAChr ɑ7 on mature human monocyte-derived macrophages and binding of recombinant trimeric RABV glycoprotein to this receptor. (A) The presence of nAChr ɑ7 on mature human MDMs, as well as the binding of ɑ-BTX and FITC-labeled rRABV-tG was visualized by confocal laser scanning microscopy. Nuclei were stained with Hoechst 33342. The white scalebar represents 500 μM. The rightmost plot shows an overlaid image of the nAChr ɑ7 staining and ɑ-BTX and rRABV-tG binding. Presence of nAChr ɑ7 was confirmed by flow cytometry (B) and this technique was also used to quantify binding of FITC-labeled rRABV-tG, shown by the mean-fluorescent intensity (MFI) of the FITC channel representing rRABV-tG binding, as well as the blocking of binding by pre-incubation with the nAChr ɑ7 specific antagonist ɑ-BTX (C). Bars represent the mean ± SEM for a total of six donors. *P* values < 0.05 were considered significant and are indicated with an asterisk (*). Non-significant comparisons were indicated with n.s.

### Binding of RABV to nAChr ɑ7 induces the cholinergic anti-inflammatory pathway in human macrophages through cytoplasmic retention of NF-κB

After confirming that rRABV-tG binds to nAChr ɑ7 on human macrophages we set out to investigate whether this binding could result in induction of the cholinergic anti-inflammatory pathway (CAP). Mature MDMs were exposed to various doses of RABV (MOI = 10, 25, 50), after which they were stimulated with LPS for six hours. Quantification of TNF-ɑ by cytometric bead assays showed that exposure to high concentrations of RABV (MOI of 50) resulted in a significant decrease in TNF-ɑ upon stimulation with LPS (Fig 3A). The decrease in TNF-ɑ production (30.1 % on average) was similar to the decrease observed with wells pre-treated with acetylcholine, a prototypical ligand of nAChr ɑ7. The lower concentrations of RABV (MOI of 10 and 25) did not induce this decrease, indicating that higher viral concentrations were required to induce the CAP. Importantly, by pre-treating macrophages with ɑ-BTX we were able to inhibit the induction of the CAP, as TNF-ɑ production was similar to the controls treated with LPS only. This confirms that observed induction of the CAP is caused by specific binding of RABV-G to nAChr ɑ7.

**Fig 3.**
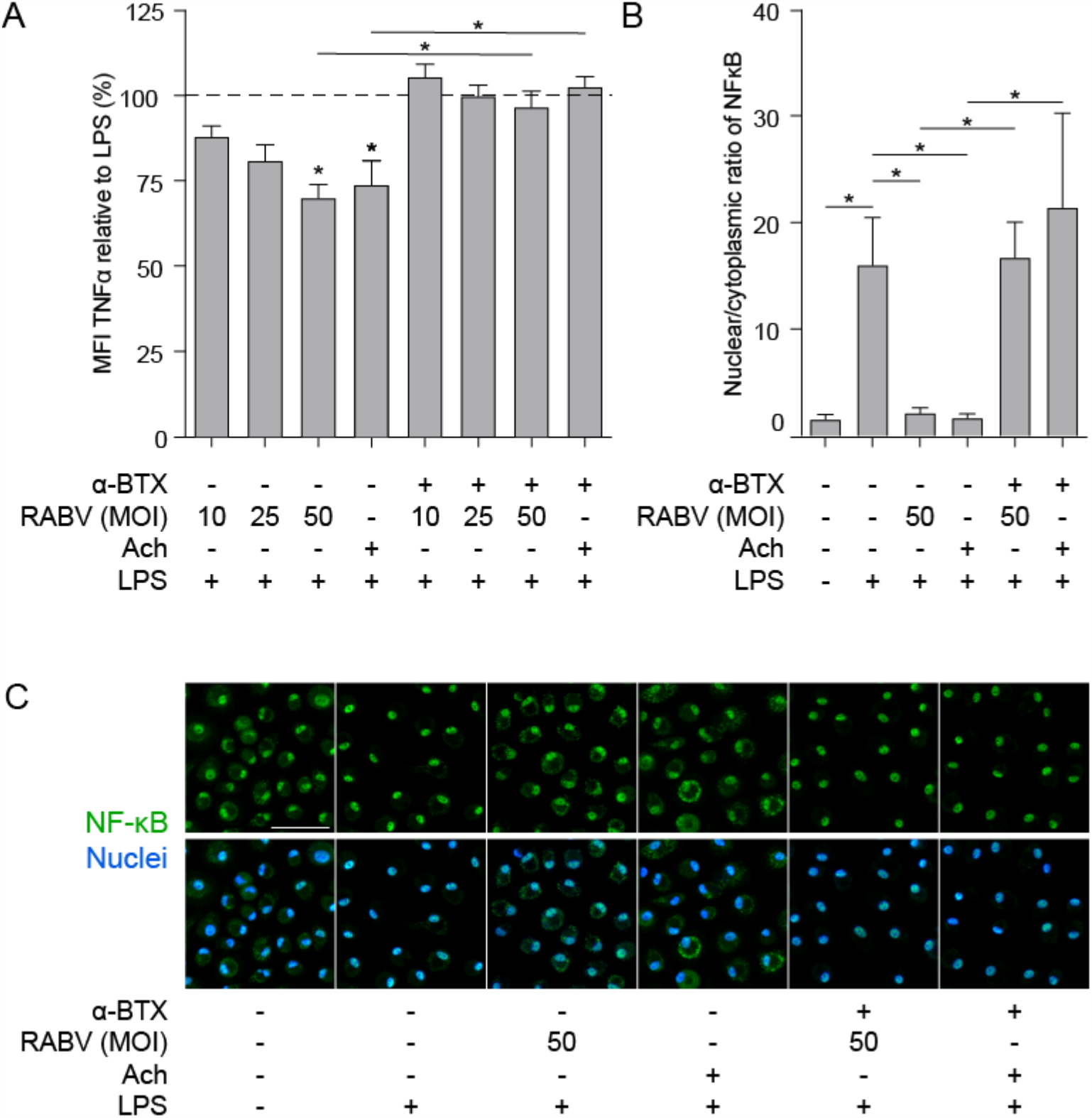
Quantification of the anti-inflammatory effect induced of RABV on human macrophages and the role of cytoplasmic retention of NF-κB. (A) TNF-ɑ production in human macrophages after six hours stimulation with LPS (100 ng/mL), with (‘+’) or without (‘-’) pre-treatment of ɑ-BTX (2 µg/mL), RABV or acetylcholine (Ach; 1 µM), or a combination. Bars represent the mean ± SEM of TNF-ɑ normalized to cells stimulated with LPS only (positive control, dotted horizontal line) for each individual donor (n = 6–9). Horizontal lines represent paired comparisons, whereas asterisks without horizontal lines indicate comparisons with the positive control (LPS only). (B) Nuclear and cytoplasmic NF-κB was quantified by image analysis in ImageJ using a batch image analysis. Bars represent the mean ± SEM of six donors, for which three high-magnification fields were analyzed per treatment. *P* values < 0.05 were considered significant and are indicated with an asterisk (*). Representative pictures of NF-κB staining are depicted in panel (C). Pictures where the NF-κB staining (green) completely overlaps with the nuclei (blue) show full nuclear translocation, whereas pictures with non-complete overlap of NF-κB and nuclei indicate cytoplasmic retention of NF-κB. The white scalebar represents 500 μM and all images were acquired at the same magnification.

Next, we aimed at investigating whether induction of the CAP by RABV also involves cytoplasmic retention of NF-κB, as is described for the prototypical ligands nicotine and acetylcholine. Macrophages were either treated with RABV (MOI of 50) or acetylcholine before challenge with LPS, after which the cells were fixed and stained with an antibody against the NF-κB p65 subunit. Nuclear and cytoplasmic NF-κB were quantified in confocal microscopy images using ImageJ batch image processing. Exposure of human MDMs to LPS led to nuclear translocation of NF-κB, hereby significantly increasing the nuclear/cytoplasmic ratio (Fig 3B and 3C). This translocation could be blocked almost completely by pre-treating macrophages with RABV or acetylcholine, causing cytoplasmic retention of NF-κB. Notably, the cytoplasmic retention caused by both RABV and acetylcholine could be blocked by pre-treating the cells with the nAChr ɑ7 specific antagonist ɑ-BTX, confirming that binding of RABV to the nAChr ɑ7 is essential for the observed NF-κB cytoplasmic retention.

### RABV-exposed macrophages suppress T cell proliferation

Macrophages play an important role in initiating adaptive immune responses by presenting antigens and producing various cytokines that either inhibit or activate T proliferation upon interaction with these cells. We investigated if the observed anti-inflammatory response of RABV on human MDMs could also affect T cell proliferation *in vitro*. To study this, we performed co-cultures of macrophages exposed to RABV (MOI of 50), and autologous T cells that had been activated with soluble ɑ-CD3 and ɑ-CD28. T cells were stained with CFSE, and after 3.5 days T cell proliferation was analyzed by flow cytometry. Proliferation of CD8^+^ T cells was significantly decreased upon co-culture with RABV-exposed macrophages (average decrease of 8.6 %) when compared to T cells cultured with control macrophages not exposed to RABV (Fig 4). Although not significant, CD4^+^ T cells showed a similar trend (average decrease of 7.8 %). This shows that exposure of human MDMs to RABV is able to suppress T cell proliferation *in vitro*.

**Fig 4.**
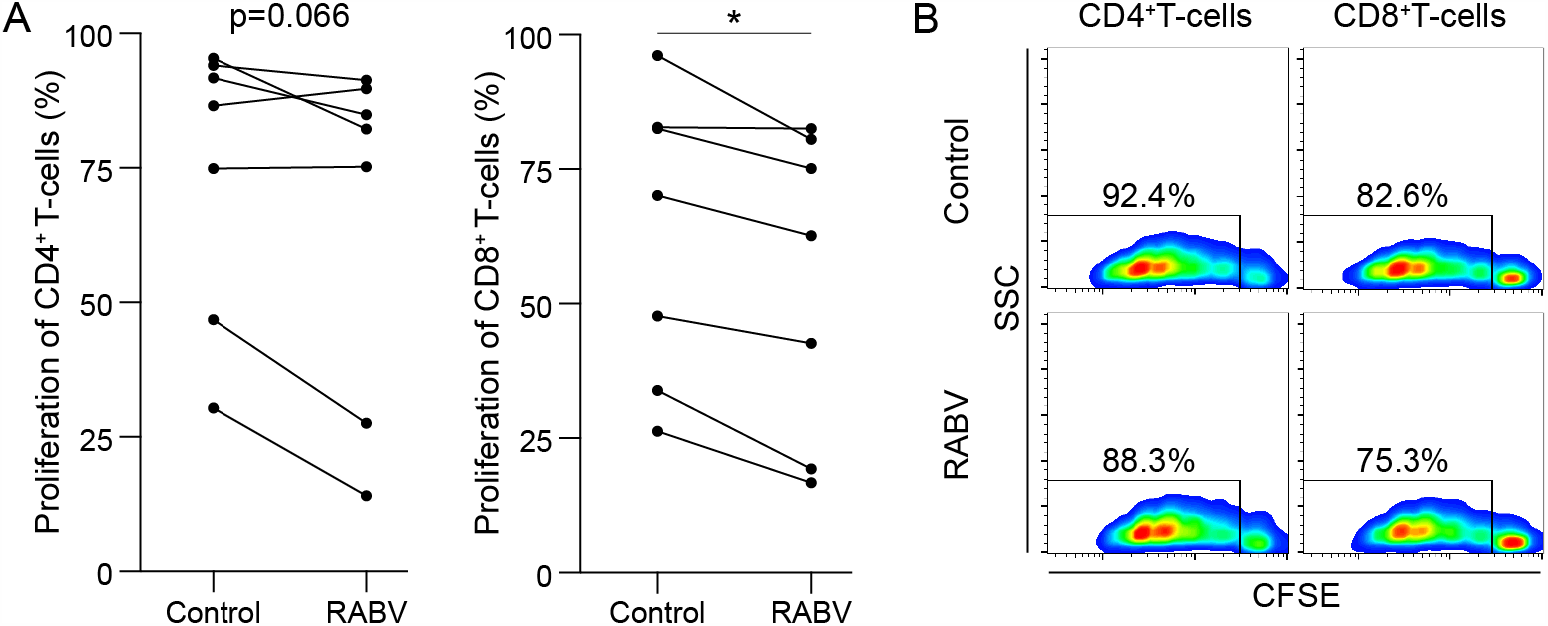
Proliferation of autologous T cells during *in vitro* co-culture with human monocyte-derived macrophages exposed to RABV. Proliferation of CFSE-stained T cells was quantified by flow cytometry after 3.5 days of co-culture with autologous MDMs previously exposed to RABV (MOI = 50). T cells cultured in the presence of macrophages that were not exposed to RABV were taken along as controls. Individual donors (n = 6) are shown in (A) and representative CFSE plots showing the fluorescent intensity and percentages of proliferation are shown in (B). *P* values < 0.05 were considered significant and are indicated with an asterisk (*).

### RABV does not polarize human macrophages towards a M1 phenotype but induces upregulation of the M2-c marker CD163

After showing that exposure to RABV induces an anti-inflammatory pathway in human MDMs, we investigated whether longer exposure (48 hours) to RABV (MOI of 10) was able to induce polarization of human MDMs. A panel of M1, M2-a and M2-c phenotypical markers was used to investigate the macrophages’ phenotype by flow cytometry; polarizing cytokine cocktails known to induce M1, M2-a and M2-c phenotypes were taken along as controls. Polarization with IFN-γ and LPS induced a typical M1 phenotype, characterized by significant upregulation of CD80, HLA-DR and PD-L1 (Fig 5). Contrary to this, the typical M2-a markers CD200R and CD206 were upregulated after polarization with IL1-4. CD163, a typical M2-c marker, was not only upregulated after polarization with IL-10, but also after exposure to RABV. Although PD-L1 was also upregulated in macrophages polarized in the presence of RABV, the upregulation was lower than observed in macrophages polarized with IL-4. All together these findings demonstrate that while RABV does not induce a typical M1 phenotype, it might steer macrophage polarization towards an M2-c phenotype, as characterized by the upregulation of CD163.

**Fig 5.**
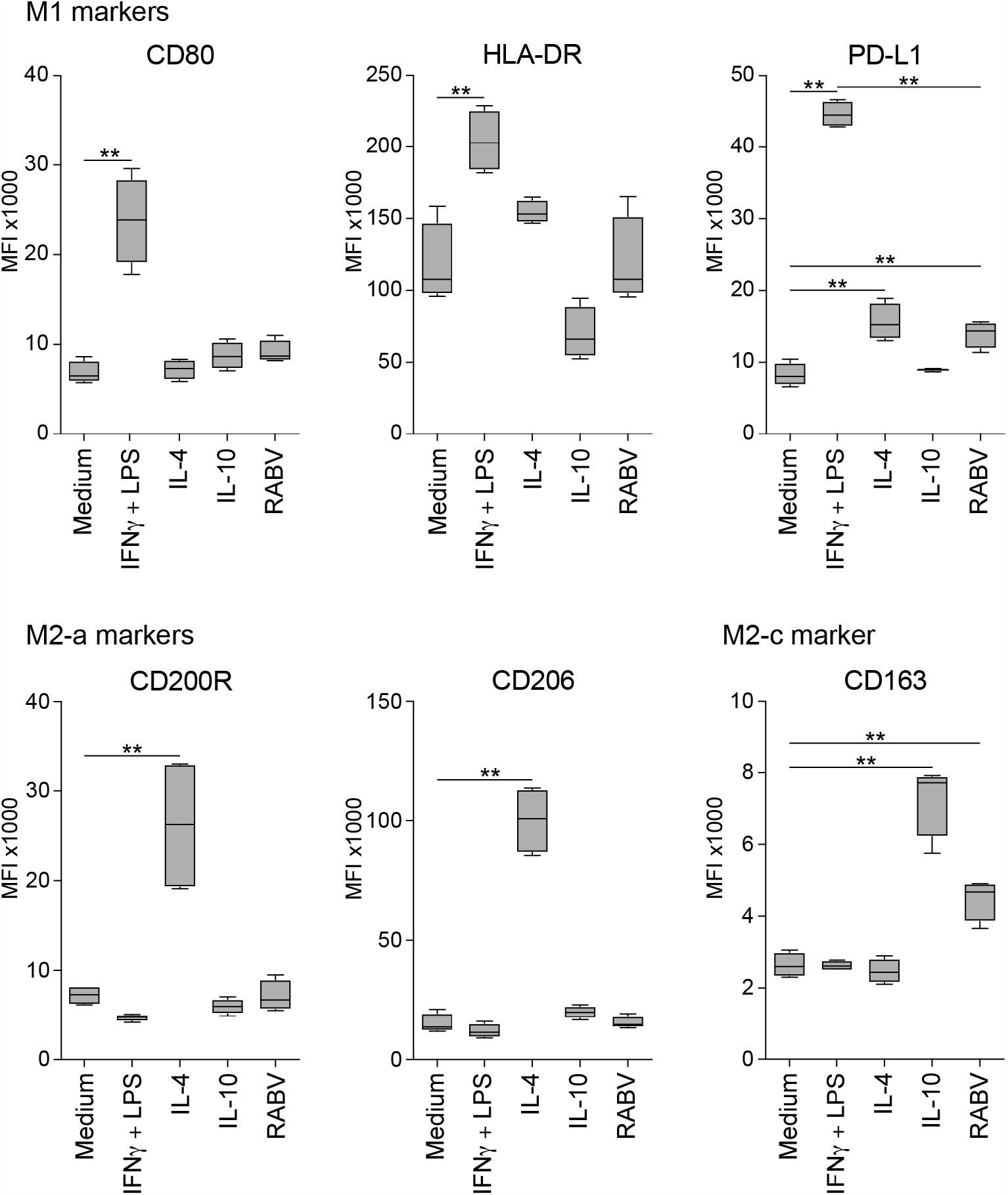
Expression levels of macrophage phenotypical markers after 48 hours of polarization with various cytokines or RABV. Panels represent typical M1 markers (CD80, HLA-DR and PD-L1), M2-a markers (CD200R and CD206) and the M2-c marker CD163. Mean fluorescent intensities (MFI) were measured by flow cytometry after 48 hours of polarization. Bars represent the mean ± SEM of six donors. Values were considered significant when *P* < 0.05 and are indicated with an asterisk (*). *P* < 0.005 are indicated with (**).

## Discussion

In this study we investigated if street RABV can induce anti-inflammatory pathways in human MDMs and if interaction between RABV and macrophages can affect T cell proliferation. Knowledge on the early interactions between RABV and local immune cells are of utmost importance in order to develop and improve efficient post-exposure treatments. We demonstrate that a street RABV strain (SHBRV) has an anti-inflammatory effect on human MDMs by inducing the CAP through binding of RABV-G to nAChr ɑ7, characterized by cytoplasmic retention of NF-κB and a decreased TNF-ɑ response upon LPS stimulation. We further show that exposure of macrophages to RABV decreases T cell proliferation *in vitro* and upregulates surface expression of CD163, a marker of the anti-inflammatory M2-c phenotype.

Before studying the immunomodulatory effects of street RABV on human MDMs, we investigated if street RABV is able to replicate in human MDMs and concluded that this is not the case (Fig 1). This lack of replication was also reported in murine bone marrow- or peritoneal-derived macrophages infected with various street RABV strains (Ray et al., 1995), indicating that primary macrophages are non-permissive to infection with street RABV strains. In contrast, primary murine macrophages did show replication of the lab-adapted CVS-11 and attenuated HEP Flury strains (Kip et al., 2017) and the recombinant matrix gene-deleted vaccine strain rRABV-ΔM (Lytle et al., 2015). This indicates that in contrast to street RABV strains, some attenuated strains are able to productively infect primary macrophages.

We next confirmed that RABV-G specifically binds to the nAChr ɑ7 on human MDMs, as binding of the recombinant trimeric rRABV-tG was decreased when cells were pre-treated with the receptor-specific antagonist ɑ-BTX. However, pre-treatment did not block binding of rRABV-tG completely, indicating that RABV-G binds to other receptors on human MDMs as well. Alternative receptors may include additional nAChr, given that the expression of a wide array of nAChr (including ɑ1, ɑ-3-6, ɑ9-10, β1-β4) have been described for human macrophages and macrophage-like cell lines (Chernyavsky et al., 2010; Galvis et al., 2006).

While major immune evasive mechanisms have been identified for the RABV nucleoprotein (N) and phosphoprotein (P), affecting RIG-I activation (Masatani et al., 2013, 2011, 2010) and the IFN signal transduction pathway (Brzózka et al., 2006, 2005; Vidy et al., 2005) or cytokine signaling (Harrison et al., 2020) respectively, related mechanisms for the surface protein RABV-G have not yet been identified. Our results show that exposure of human MDMs to RABV leads to a decreased TNF-ɑ response upon LPS challenge, caused by cytoplasmic retention of NF-κB. TNF-ɑ levels and NF-κB nuclear translocation was completely restored when MDMs were pre-treated with the nAChr ɑ7-specific antagonist ɑ-BTX, showing that the observed anti-inflammatory effects of RABV were caused by binding of the RABV-G to nAChr ɑ7. While multiple molecules have been found to induce the CAP in macrophages, including GTS-21 (van Westerloo et al., 2006), CAP55 (Saeed et al., 2005) and PNU-282987 (Dash et al., 2016), to our knowledge this is the first study showing that a viral protein is able to induce this pathway. The CAP might have additional effects on RABV pathogenesis and disease development, as was shown that the nAChr ɑ7 antagonist GTS-21 attenuates the cytokine response in monocytes after stimulation with ligands for Toll-Like Receptor 2 (TLR2), TLR3, TLR4, TLR9 and RAGE. Furthermore, downregulating the inflammatory response was also observed in microglia (Egea et al., 2015), which are resident CNS macrophages that also express nAChr-ɑ7 (Shytle et al., 2004).

Given the importance of the NF-κB signaling pathway in the initiation of an immune response, multiple viruses (including Borna disease virus, Epstein-Barr virus, Hantaan virus, Hepatitis C virus, Poliovirus, Varicella-zoster virus and West Nile virus, as extensively reviewed (Rahman and McFadden, 2011)) have acquired mechanisms to inhibit NF-κB activation. We showed that RABV induced cytoplasmic retention of NF-κB *in vitro*, which is described as an essential step in activation of the CAP (Wang et al., 2004). For RABV it is known that the matrix (M) protein is able to inhibit RelAP43 activation, a splice variant of the NF-κB subunit RelA, leading to decreased expression of various innate immune genes (Benoit et al., 2017; Khalifa et al., 2016; Luco et al., 2012). Similar effects have not yet been described for the other RABV proteins and our study is the first to show that RABV-G is able to specifically inhibit activation of the NF-κB signaling pathway. Contrary to our findings, microglia-like cell lines infected with the attenuated strain CVS-11 showed a strong activation of NF-κB (Nakamichi et al., 2005). However, productive infection was observed in the cell lines used in that study.

Macrophages are APCs and potent cytokine producing cells, and therefore suppressed macrophage functioning can lead to decreased T cell activation. T cell activation requires three signals: T cell receptor (TCR) binding to antigens presented on major histocompatibility complexes (MHC) of APCs (signal 1), binding of costimulatory molecules (signal 2), and activation by cytokines. It has been proposed that IL-1 can provide the third signal for CD4^+^ cells (Curtsinger and Mescher, 2010; Huber et al., 1998), resulting in T cell activation. Given that the inhibition of IL-1 had been described after induction of the CAP in macrophages (Sugano et al., 1998; Takahashi et al., 2006) the observed decrease in T cell proliferation might be caused by a decreased activation of CD4^+^ T cells. As a consequence, decreased cytokine production by CD4^+^ cells, and especially IL-2, can be a probable explanation for the significant decrease in CD8^+^ T cell proliferation.

Viral antigens can be sensed by macrophages and are capable of steering macrophage polarization in a certain direction (Brancato and Albina, 2011; Daley et al., 2010); the classically activated M1, or pro-inflammatory macrophage, and the M2 macrophage, also known as alternatively activated or anti-inflammatory macrophages (Koh and Dipietro, 2019; Mosser and Edwards, 2008). Generally, macrophages polarize towards an M1 phenotype after recognition of intracellular pathogens, including viruses. We showed that exposure of human MDMs to a street RABV strain for 2 days significantly upregulated the M2-c marker CD163, while the typical M1-markers CD80 and HLA-DR remained unchanged. While more in-depth investigation on surface marker expression and macrophage functioning is necessary, the results hint at the ability of street RABV to shift macrophage polarization towards an M2-c phenotype. Studies on the effects of street RABV strains on polarization of primary macrophages are lacking, but studies using attenuated viruses CVS-11 and HEP show an opposite effect. Upregulated gene expression of iNOS and nitric oxide (NO) expression, a key characteristic of M1 macrophages, in the murine macrophage-like cell like RAW264 (Nakamichi et al., 2004), and the attenuated SPBN-GAS was able to shift the polarization pattern of tumor-associated macrophages (TAMs), that normally resemble a M2-phenotype, towards an M1-phenotype in glioma-bearing mice (Bongiorno et al., 2017).

CD163 is a scavenger receptor within the cysteine-rich family and is a marker of M2-c anti-inflammatory macrophages. Increased CD163 expression on macrophages has been reported during viral hepatitis (Hiraoka et al., 2005), and infection of pigs with porcine reproductive and respiratory syndrome virus (Patton et al., 2009) and African swine fever virus (Sánchez-Torres et al., 2003). Furthermore, an increase in CD163^+^ macrophages was found in individuals infected with human immunodeficiency virus type 1 (Fischer-Smith et al., 2008), as well as in rhesus macaques infected with simian immunodeficiency virus (Yearley et al., 2007). Interestingly, the latter study showed a negative correlation between CD163^+^ macrophages and inflammatory infiltration in areas infected, indicating that CD163^+^ macrophages serve an anti-inflammatory or immunosuppressive role. In addition, CD163^+^ TAMs were found to suppress T cell proliferation in several disease models (Han et al., 2016; Lepique et al., 2009; Lievense et al., 2016; Oishi et al., 2016).

In summary, our results show for the first time that a viral protein, the RABV-G of a street RABV strain, is able to induce the CAP in human MDMs. We show that this anti-inflammatory pathway is induced by binding of RABV-G to nAChr ɑ7, using the specific antagonist ɑ-BTX, leading to cytoplasmic retention of NF-κB. Besides the decreased inflammatory response upon challenge with LPS we found that exposing human MDMs to RABV leads to suppression of T cell proliferation *in vitro*. We also show that exposure of human MDMs to RABV does not induce a typical, inflammatory M1 phenotype, but instead a significant upregulation of CD163, marker of a M2-c anti-inflammatory phenotype. Given that the absence of a strong local innate immune response is beneficial for the virus, polarizing resident and infiltrating macrophages towards an anti-inflammatory M2-c phenotype might be another mechanism of RABV to evade the immune system.

Future studies should focus on further understanding of the observed lack of adaptive immune response. Additional *in vitro* experiments in which the effects of “RABV-polarized” macrophages on T and B cells are investigated will allow evaluation of the role of macrophage suppression on the lack of neutralizing antibodies in RABV patients. *In vivo* experiments using targeted transgenic mice will allow reveal the role of macrophage suppression on the complete course of disease. Altogether, thorough insights into the different mechanisms that RABV uses to suppress the immune system are essential for the development of new and improved PEP and treatment options in RABV infection.

## Acknowledgements

The authors thank Janneke Hendriks for her enthusiasm during the Master project. Georgina Arron and Petra van den Doel are acknowledged for excellent technical support and Rudi Hendriks for scientific discussions.

## Notes

### Competing Interest Statement

The authors have declared no competing interest.

## References

Benoit, B., Sonthonnax, F., Duchatea, M., Khalifa, Y. Ben, Larrous, F., Eun, H., Hourdel, V., Matondo, M., Chamot-Rooke, J., Grailhe, R., Bourhy, H., 2017. Regulation of NF-kB by the p105-ABIN2-TPL2 complex and RelAp43 during rabies virus infection. PLoS Pathog. 1–24.

Bongiorno, E.K., Garcia, S.A., Sauma, S., Hooper, D.C., 2017. Type 1 Immune Mechanisms Driven by the Response to Infection with Attenuated Rabies Virus Result in Changes in the Immune Bias of the Tumor Microenvironment and Necrosis of Mouse GL261 Brain Tumors. J. Immunol. 198, 4513–4523. doi:10.4049/jimmunol.1601444

Brancato, S.K., Albina, J.E., 2011. Wound Macrophages as Key Regulators of Repair. Am. J. Pathol. 178, 19–25. doi:10.1016/j.ajpath.2010.08.003

Brzózka, K., Finke, S., Conzelmann, K.-K., 2006. Inhibition of Interferon Signaling by Rabies Virus Phosphoprotein P: Activation-Dependent Binding of STAT1 and STAT2. J. Virol. 80, 2675–2683. doi:10.1128/jvi.80.6.2675-2683.2006

Brzózka, K., Finke, S., Conzelmann, K.-K., 2005. Identification of the rabies virus alpha/beta interferon antagonist: phosphoprotein P interferes with phosphorylation of Interferon Regulatory factor 3. J. Virol. 79, 7673–7681. doi:10.1128/JVI.79.12.7673

Charlton, K.M., Casey, G.A., 1981. Experimental rabies in skunks: persistence of virus in denervated muscle at the inoculation site. Can. J. Comp. Med. 45, 357–362.

Charlton, K.M., Nadin-Davis, S., Casey, G.A., Wandeler, A.I., 1997. The long incubation period in rabies: Delayed progression of infection in muscle at the site of exposure. Acta Neuropathol. 94, 73–77. doi:10.1007/s004010050674

Chernyavsky, A.I., Arredondo, J., Skok, M., Grando, S.A., 2010. Auto/paracrine control of inflammatory cytokines by acetylcholine in macrophage-like U937 cells through nicotinic receptors. Int. Immunopharmacol. 10, 308–315. doi:10.1016/j.intimp.2009.12.001

Curtsinger, J.M., Mescher, M.F., 2010. Inflammatory cytokines as a third signal for T cell activation. Curr. Opin. Immunol. 22, 333–340. doi:10.1038/jid.2014.371

Daley, J.M., Brancato, S.K., Thomay, A.A., Reichner, J.S., Albina, J.E., 2010. The phenotype of murine wound macrophages. J. Leukoc. Biol. 87, 59–67. doi:10.1189/jlb.0409236

Dash, P.K., Zhao, J., Kobori, N., Redell, J.B., Hylin, M.J., Hood, K.N., Moore, A.N., 2016. Activation of alpha 7 cholinergic nicotinic receptors reduce blood–brain barrier permeability following experimental traumatic brain injury. J. Neurosci. 36, 2809–2818. doi:10.1523/JNEUROSCI.3197-15.2016

Dietzschold, B., Morimoto, K., Hooper, D.C., Smith, J.S., Rupprecht, C.E., Koprowski, H., 2000. Genotypic and phenotypic diversity of rabies virus variants involved in human rabies: implications for postexposure prophylaxis. J. Hum. Virol. 3, 50–57.

Egea, J., Buendia, I., Parada, E., Navarro, E., León, R., Lopez, M.G., 2015. Anti-inflammatory role of microglial alpha7 nAChRs and its role in neuroprotection. Biochem. Pharmacol. 97, 463–472. doi:10.1016/j.bcp.2015.07.032

Fischer-Smith, T., Tedaldi, E.M., Rappaport, J., 2008. CD163/CD16 coexpression by circulating monocytes/macrophages in HIV: Potential biomarkers for HIV infection and AIDS progression. AIDS Res. Hum. Retroviruses 24, 417–421. doi:10.1089/aid.2007.0193

Galvis, G., Lips, K.S., Kummer, W., 2006. Expression of nicotinic acetylcholine receptors on murine alveolar macrophages. J. Mol. Neurosci. 30, 107. doi:10.1385/JMN/30:1-2:107

Hampson, K., Coudeville, L., Lembo, T., Sambo, M., Kieffer, A., Attlan, M., Barrat, J., Blanton, J.D., Briggs, D.J., Cleaveland, S., Costa, P., Freuling, C.M., Hiby, E., Knopf, L., Leanes, F., Meslin, F.X., Metlin, A., Miranda, M.E., Müller, T., Nel, L.H., Recuenco, S., Rupprecht, C.E., Schumacher, C., Taylor, L., Vigilato, M.A.N., Zinsstag, J., Dushoff, J., 2015. Estimating the Global Burden of Endemic Canine Rabies. PLoS Negl. Trop. Dis. 9, 1–20. doi:10.1371/journal.pntd.0003709

Han, Q., Shi, H., Liu, F., 2016. CD163 + M2-type tumor-associated macrophage support the suppression of tumor-infiltrating T cells in osteosarcoma. Int. Immunopharmacol. 34, 101– 106. doi:10.1016/j.intimp.2016.01.023

Harrison, A.R., Lieu, K.G., Larrous, F., Ito, N., Bourhy, H., Moseley, G.W., 2020. Lyssavirus P-protein selectively targets STAT3-STAT1 heterodimers to modulate cytokine signalling. PLoS Pathog. 1–23. doi:10.1371/journal.ppat.1008767

Hiraoka, A., Horiike, N., Fazle Akbar, S.M., Michitaka, K., Matsuyama, T., Onji, M., 2005. Expression of CD163 in the liver of patients with viral hepatitis. Pathol. Res. Pract. 201, 379–384. doi:10.1016/j.prp.2004.10.006

Huber, M., Beuscher, H.U., Rohwer, P., Kurrle, R., Röllinghoff, M., Lohoff, M., 1998. Costimulation via TCR and IL-1 receptor reveals a novel IL-1ɑ-mediated autocrine pathway of TH2 cell proliferation. J. Immunol. 160, 4242–4247.

Kasempimolporn, S., Hemachudha, T., Khawplod, P., Manatsathit, S., 1991. Human immune response to rabies nucleocapsid and glycoprotein antigens. Clin. Exp. Immunol. 84, 195– 199. doi:10.1111/j.1365-2249.1991.tb08148.x

Katz, I.S.S., Guedes, F., Fernandes, E.R., Dos Ramos Silva, S., 2017. Immunological aspects of rabies: a literature review. Arch. Virol. 162, 1–18. doi:10.1007/s00705-017-3484-0

Khalifa, Y. Ben, Luco, S., Besson, B., Sonthonnax, F., Archambaud, M., Grimes, J.M., Larrous, F., Bourhy, H., 2016. The matrix protein of rabies virus binds to RelAp43 to modulate NF-κB-dependent gene expression related to innate immunity. Sci. Rep. 6, 1–13. doi:10.1038/srep39420

Kim, S.-S., Ye, C., Kumar, P., Chiu, I., Subramanya, S., Wu, H., Shankar, P., Manjunath, N., 2010. Targeted Delivery of siRNA to Macrophages for Anti-inflammatory Treatment. Mol. Ther. 18, 993–1001. doi:10.1038/mt.2010.27

Kip, E., Nazé, F., Suin, V., Vanden Berghe, T., Francart, A., Lamoral, S., Vandenabeele, P., Beyaert, R., Van Gucht, S., Kalai, M., 2017. Impact of caspase-1/11, −3, −7, or IL-1β/IL-18 deficiency on rabies virus-induced macrophage cell death and onset of disease. Cell Death Discov. 3, 17012. doi:10.1038/cddiscovery.2017.12

Koh, T.J., Dipietro, L.A., 2019. Inflammation and wound healing?: the role of the macrophage. Expert Rev. Mol. Med. 13, 1–12. doi:10.1017/S1462399411001943

Koraka, P., Bosch, B.J., Cox, M., Chubet, R., Amerongen, G.van, Lövgren-Bengtsson, K., Martina, B.E.E., Roose, J., Rottier, P.J.M., Osterhaus, A.D.M.E., 2014. A recombinant rabies vaccine expressing the trimeric form of the glycoprotein confers enhanced immunogenicity and protection in outbred mice. Vaccine 32, 4644–4650. doi:10.1016/j.vaccine.2014.06.058

Lepique, A.P., Daghastanli, K.R.P., Cuccovia, I., Villa, L.L., 2009. HPV16 tumor associated macrophages suppress antitumor T cell responses. Clin. Cancer Res. 15, 4391–4400. doi:10.1158/1078-0432.CCR-09-0489

Lievense, L.A., Cornelissen, R., Bezemer, K., Kaijen-Lambers, M.E.H., Hegmans, J.P.J.J., Aerts, J.G.J.V., 2016. Pleural effusion of patients with malignant mesothelioma induces macrophage-mediated T Cell suppression. J. Thorac. Oncol. 11, 1755–1764. doi:10.1016/j.jtho.2016.06.021

Luco, S., Delmas, O., Vidalain, P.O., Tangy, F., Weil, R., Bourhy, H., 2012. RelAp43, a Member of the NF-κB Family Involved in Innate Immune Response against Lyssavirus Infection. PLoS Pathog. 8. doi:10.1371/journal.ppat.1003060

Lytle, A.G., Shen, S., McGettigan, J.P., 2015. Lymph node but not intradermal injection site macrophages are critical for germinal center formation and antibody responses to rabies vaccination. J. Virol. 89, 2842–8. doi:10.1128/JVI.03409-14

Masatani, T., Ito, N., Ito, Y., Nakagawa, K., Abe, M., Yamaoka, S., Okadera, K., Sugiyama, M., 2013. Importance of rabies virus nucleoprotein in viral evasion of interferon response in the brain. Microbiol. Immunol. 57, 511–517. doi:10.1111/1348-0421.12058

Masatani, T., Ito, N., Shimizu, K., Ito, Y., Nakagawa, K., Abe, M., Yamaoka, S., Sugiyama, M., 2011. Amino acids at positions 273 and 394 in rabies virus nucleoprotein are important for both evasion of host RIG-I-mediated antiviral response and pathogenicity. Virus Res. 155, 168–174. doi:10.1016/j.virusres.2010.09.016

Masatani, T., Ito, N., Shimizu, K., Ito, Y., Nakagawa, K., Sawaki, Y., Koyama, H., Sugiyama, M., 2010. Rabies Virus Nucleoprotein Functions To Evade Activation of the RIG-I-Mediated Antiviral Response. J. Virol. 84, 4002–4012. doi:10.1128/jvi.02220-09

Mosser, D.M., Edwards, J.P., 2008. Exploring the full spectrum of macrophage activation. Nat. Rev. Immunol. 8, 958–969. doi:10.1038/nri2448

Murphy, A.M., Bauer, S.P., 1974. Early street rabies virus infection in striated muscle and later progression to the central nervous system. Intervirology 3, 256–268.

Nakamichi, K., Inoue, S., Takasaki, T., Morimoto, K., Kurane, I., 2004. Rabies Virus Stimulates Nitric Oxide Production and CXC Chemokine Ligand 10 Expression in Macrophages through Activation of Extracellular Signal-Regulated Kinases 1 and 2. J. Virol. 78, 9376– 9388. doi:10.1128/JVI.78.17.9376

Nakamichi, K., Saiki, M., Sawada, M., Yamamuro, Y., Morimoto, K., Kurane, I., Takayama-ito, M., 2005. Rabies Virus-Induced Activation of Mitogen-Activated Protein Kinase and NF-κ B Signaling Pathways Regulates Expression of CXC and CC Chemokine Ligands in Microglia. J. Virol. 79, 11801–11812. doi:10.1128/JVI.79.18.11801

Noah, D.L., Drenzek, C.L., Smith, J.S., Krebs, J.W., Orciari, L., Shaddock, J., Sanderlin, D., Whitfield, S., Fekadu, M., Olson, J.G., Rupprecht, C.E., Childs, J.E., 1998. Epidemiology of human rabies in the United States, 1980 to 1996. Ann. Intern. Med. 128, 922–930. doi:10.7326/0003-4819-128-11-199806010-00012

Oishi, S., Takano, R., Tamura, S., Tani, S., Iwaizumi, M., Hamaya, Y., Takagaki, K., Nagata, T., Seto, S., Horii, T., Osawa, S., Furuta, T., Miyajima, H., Sugimoto, K., 2016. M2 polarization of murine peritoneal macrophages induces regulatory cytokine production and suppresses T-cell proliferation. Immunology 149, 320–328. doi:10.1111/imm.12647

Patton, J.B., Rowland, R.R., Yoo, D., Chang, K.O., 2009. Modulation of CD163 receptor expression and replication of porcine reproductive and respiratory syndrome virus in porcine macrophages. Virus Res. 140, 161–171. doi:10.1016/j.virusres.2008.12.002

Piazzon, M.C., Savelkoul, H.F.J., Pietretti, D., Wiegertjes, G.F., Forlenza, M., 2015. Carp Il10 Has Anti-Inflammatory Activities on Phagocytes, Promotes Proliferation of Memory T Cells, and Regulates B Cell Differentiation and Antibody Secretion. J. Immunol. 194, 187– 199. doi:10.4049/jimmunol.1402093

Rahman, M.M., McFadden, G., 2011. Modulation of NF-κB signalling by microbial pathogens. Nat. Rev. Microbiol. 9, 291–306. doi:10.1038/nrmicro2539

Ray, N.B., Ewalt, L.C., Lodmell, D.L., 1995. Rabies virus replication in primary murine bone marrow macrophages and in human and murine macrophage-like cell lines: implications for viral persistence. J. Virol. 69, 764–772.

Reed, L.J., Muench, H., 1938. A simple method of estimating fifty per cent endpoints. Am. J. Hyg. 27, 546–558. doi:10.7723/antiochreview.72.3.0546

Saeed, R.W., Varma, S., Peng-Nemeroff, T., Sherry, B., Balakhaneh, D., Huston, J., Tracey, K.J., Al-Abed, Y., Metz, C.N., 2005. Cholinergic stimulation blocks endothelial cell activation and leukocyte recruitment during inflammation. J. Exp. Med. 201, 1113–1123. doi:10.1084/jem.20040463

Sánchez-Torres, C., Gómez-Puertas, P., Gómez-Del-Moral, M., Alonso, F., Escribano, J.M., Ezquerra, A., Domínguez, J., 2003. Expression of porcine CD163 on monocytes/macrophages correlates with permissiveness to African swine fever infection. Arch. Virol. 148, 2307–2323. doi:10.1007/s00705-003-0188-4

Sang, Y., Miller, L.C., Blecha, F., 2015. Macrophage Polarization in Virus-Host Interactions. J. Clin. Cell. Immunol. 06. doi:10.4172/2155-9899.1000311

Schnell, M.J., McGettigan, J.P., Wirblich, C., Papaneri, A., 2010. The cell biology of rabies virus: Using stealth to reach the brain. Nat. Rev. Microbiol. 8, 51–61. doi:10.1038/nrmicro2260

Scott, T.P., Nel, L.H., 2016. Subversion of the immune response by Rabies Virus. Viruses 8, 1– 26. doi:10.3390/v8080231

Shytle, R.D., Mori, T., Townsend, K., Vendrame, M., Sun, N., Zeng, J., Ehrhart, J., Silver, A.A., Sanberg, P.R., Tan, J., 2004. Cholinergic modulation of microglial activation by ɑ7 nicotinic receptors. J. Neurochem. 89, 337–343. doi:10.1046/j.1471-4159.2004.02347.x

Stout, R.D., Jiang, C., Matta, B., Tietzel, I., Watkins, S.K., Suttles, J., 2005. Macrophages Sequentially Change Their Functional Phenotype in Response to Changes in Microenvironmental Influences. J. Immunol. 175, 342–349. doi:10.4049/jimmunol.175.1.342

Sugano, N., Shimada, K., Ito, K., Murai, S., 1998. Nicotine inhibits the production of inflammatory mediators in U937 cells through modulation of Nuclear Factor-kB activation. Biochem. Biophys. Res. Commun. 252, 25–28. doi:10.1006/bbrc.1998.9599

Takahashi, H.K., Iwagaki, H., Hamano, R., Yoshino, T., Tanaka, N., Nishibori, M., 2006. Effect of nicotine on IL-18-initiated immune response in human monocytes. J. Leukoc. Biol. 80, 1388–1394. doi:10.1189/jlb.0406236

Turner, G.S., Ballard, R., 1976. Interaction of mouse peritoneal macrophages with fixed rabies virus in vivo and in vitro. J. Gen. Virol. 30, 223–231. doi:10.1099/0022-1317-30-2-223

van Westerloo, D.J., Giebelen, I.A., Florquin, S., Bruno, M.J., LaRosa, G.J., Ulloa, L., Tracey, K.J., van der Poll, T., 2006. The Vagus Nerve and Nicotinic Receptors Modulate Experimental Pancreatitis Severity in Mice. Gastroenterology 130, 1822–1830. doi:10.1053/j.gastro.2006.02.022

Vidy, A., Chelbi-Alix, M., Blondel, D., 2005. Rabies Virus P Protein Interacts with STAT1 and Inhibits Interferon Signal Transduction Pathways. J. Virol. 79, 14411–14420. doi:10.1128/jvi.79.22.14411-14420.2005

Wang, Hong, Liao, H., Ochani, M., Justiniani, M., Lin, X., Yang, L., Al-Abed, Y., Wang, Haichao, Metz, C., Miller, E.J., Tracey, K.J., Ulloa, L., 2004. Cholinergic agonists inhibit HMGB1 release and improve survival in experimental sepsis. Nat. Med. 10, 1216–1221. doi:10.1038/nm1124

Wang, Hong, Yu, M., Ochani, M., Amella, C.A., Tanovic, M., Susarla, S., Li, J.H., Wang, Haichao, Yang, H., Ulloa, L., Al-Abed, Y., Czura, C.J., Tracey, K.J., 2003. Nicotinic acetylcholine receptor ɑ7 subunit is an essential regulator of inflammation. Nature 421, 384–388. doi:10.1038/nature01339

WHO, 2018. Expert consultation on rabies - Third report, World Health Organization - Technical Report Series. Geneva.

Yamaoka, S., Ito, N., Ohka, S., Kaneda, S., Nakamura, H., Agari, T., Masatani, T., Nakagawa, K., Okada, K., Okadera, K., Mitake, H., Fujii, T., Sugiyama, M., 2013. Involvement of the rabies virus phosphoprotein gene in neuroinvasiveness. J. Virol. 87, 12327–12338. doi:10.1128/JVI.02132-13

Yearley, J.H., Pearson, C., Shannon, R.P., Mansfield, K.G., 2007. Phenotypic variation in myocardial macrophage populations suggests a role for macrophage activation in SIV-associated cardiac disease. AIDS Res. Hum. Retroviruses 23, 515–524. doi:10.1089/aid.2006.0211

